# Intrinsically regulated learning is modulated by synaptic dopamine availability

**DOI:** 10.1101/318576

**Authors:** Pablo Ripollés, Laura Ferreri, Ernest Mas-Herrero, Helena Alicart, Alba Gómez-Andrés, Josep Marco-Pallarés, Rosa Antonijoan, Toemme Noesselt, Marta Valle, Jordi Riba, Antoni Rodriguez-Fornells

## Abstract

We recently provided evidence that an intrinsic reward-related signal—triggered by successful learning in absence of *any external feedback*—modulated the entrance of new information into long-term memory via the activation of the dopaminergic midbrain, hippocampus, and ventral striatum (the SN/VTA-Hippocampal loop; Ripollés et al., 2016). Here, we used a double-blind, within-subject randomized pharmacological intervention to test whether this learning process is indeed dopamine-dependent. A group of healthy individuals completed three behavioural sessions of our learning task after the intake of different pharmacological treatments: a dopaminergic precursor, a dopamine receptor antagonist or a placebo. Results show that the pharmacological intervention bidirectionally modulated behavioral measures of both learning and pleasantness, inducing memory benefits after 24 hours only for those participants with a high sensitivity to reward. These results provide causal evidence for a dopamine-dependent mechanism instrumental in intrinsically regulated learning, and further suggest that subject-specific dopamine sensitivity drastically alters learning success.

## INTRODUCTION

Growing evidence both from animal and human studies support the notion that midbrain dopaminergic neurons of the substantia nigra/ventral tegmental area complex (SN/VTA), along with the the ventral striatum (VS) and the hippocampus (HP), form a functional loop (the SN/VTA-HP loop) in the service of learning and memory (Lisman and Grace, 2005; Goto and Grace, 2005; Lisman et al., 2011; Shohamy and Adcock, 2010; Kaminski et al., 2018). In the downward arm of the circuit, signals are sent from the HP to the SN/VTA through the VS, which is thought to integrate affective, motivational, and goal-directed information into the loop (Lisman and Grace, 2005; Goto and Grace, 2005). In the upward arm of the loop, dopamine is released from the SN/VTA back into the HP, which in turn enhances memory formation and learning through long term potentiation (LTP) processes (Lisman et al., 2011; Lisman and Grace, 2005; Shohamy and Adcock, 2010). Within this loop, dopamine plays a critical role, as its release promotes the creation of stable memories by allowing LTP to persist over time (Bethus et al., 2010; Frey et al., 1990; Hansen and Manahan-Vaughan, 2014; Huang and Kandel, 1995; McNamara et al., 2014; Rossato et al., 2009).

In this vein, fMRI research in humans has consistently shown that both explicit (Adcock et al., 2006; Wittmann et al., 2005; Wolosin et al., 2012; Callan et al., 2008) and implicit *reward* (Ripollés et al., 2016) can promote the storage of new information into long-term memory through the activation of the SN/VTA-HP loop (see Fig. 8 in Ripollés et al., 2016). However, although fMRI activity within the SN/VTA is usually associated with the release of dopamine (Duzel et al., 2009; Ferenczi et al., 2016; Knutson and Gibbs, 2007; Salimpoor et al., 2011; Schott et al., 2008), neuroimaging studies can only provide *indirect* evidence of the actual involvement of the dopaminergic mesolimbic system in learning and memory processes. In order to proof that a dopamine-dependent mechanism plays a critical role in this process, one avenue to pursue is to directly manipulate the levels of dopamine in the human brain through pharmacological interventions. Several studies have shown that the intake of dexamphetamine and methylphenidate (which block dopaminergic and adrenergic re-uptake; Breitenstein et al., 2004; Whiting et al., 2007; Whiting et al., 2008; Linssen et al., 2014) and specially, levodopa—a dopamine precursor—can enhance memory and learning in both healthy (Shellshear et al., 2015; Bunzeck, et al., 2014; Chowdury et al., 2012;Knecht et al., 2004) and clinical populations (Berthier et al., 2011).

We recently provided behavioural, functional and physiological evidence by means of fMRI and skin conductance response, showing that an intrinsic reward-related signal— triggered by successful learning in absence of *any external feedback or explicit reward*— modulated the entrance of new information into long-term memory via the activation of the SN/VTA-HP loop (Ripollés et al., 2016). Here, we used a double-blind, within-subject randomized pharmacological intervention to directly assess the hypothesis that synaptic dopamine availability plays a causal role in this learning process. A group of 29 individuals were asked to perform a learning task (that mimics our capacity to learn the meaning of new-words presented in verbal contexts; Ripollés et al., 2016, 2017 and 2014; Mestres-Missé et al., 2007) after the intake of three different pharmacological treatments: a dopaminergic precursor (levodopa, 100 mg + carbidopa, 25 mg), a dopamine receptor antagonist (risperidone, 2mg), or a placebo (lactose). We predicted that behavioral measures of both learning and reward should respectively increase and decrease under levodopa and risperidone, thus modulating the memory benefits for the learned words after the consolidation period (24 hours).

## Results

Twenty-nine healthy participants completed a behavioural version of our word-learning task (see Materials and Methods), in which the meaning of a new-word could be learned from the context provided by two sentences built with an increasing degree of contextual constraint (Mestres-Missé et al., 2010). Only half of the pairs of sentences disambiguated multiple meanings, allowing the encoding of a congruent meaning of the new-word during its second presentation (M+ condition). For the other pairs, the new-word was not associated with a congruent meaning across the sentences, and could not be learned (M-condition). This condition, as in our previous study (Ripollés et al., 2016), was included to control for possible confounds related to novelty, attention and task difficulty (Guitart-Masip et al., 2010; Bunzeck and Duzel, 2006; Boehler et al., 2011). At the end of each learning trial (i.e., after the second sentence for a particular new-word appeared) participants first provided a confidence rating (a subjective evaluation of their performance) and then rated their emotions with respect to arousal and pleasantness. After approximately 24 hours (no drug intake occurred during the second day of testing), participants completed a recognition test to assess their learning (chance level was 25%; see Materials and Methods). Three participants were excluded from the analyses (see Materials and Methods) and thus the final sample was reduced to 26 individuals (17 women, mean age=22.27 ± 3.69).

We first assessed whether our participants’ performance under the placebo condition replicated our previous results. Participants ascribed correct meaning to 60 ± 10 % of new-words from the M+ condition during the encoding phase. In 61 ± 15% of the M-trials, participants correctly indicated an absence of coherent meaning. After 24 hours, participants still recognized the correct meaning of 65 ± 17 % of learned new-words during the encoding phase [significantly above 25% chance level, *t*(25)=12.28, p<0.001, d=2.33; Bayes Factor-BF_10_- equal to 1.9e+9] and correctly indicated that 41 ± 22 % of M-new-words identified during the encoding phase had no meaning ascribed [significantly above 25% chance level, *t*(25)=3.70, p<0.001, d=0.70; BF_10_=35.38].

In order to compare this performance with our previous results (24 participants from Exp. 3 in Ripollés et al., 2016), we submitted the learning scores to a mixed repeated measures ANOVA with Condition (M+,M-) as a within-subjects variable and Group (Pharmacological Group, Exp. 3 in Ripollés et al., 2016) as a between subjects variable. No significant effect of Group [Learning Day 1: *F*(1,48)=0.246, p=0.622, partial η2=0.005, BF_Inclusion_=0.297; Recognition Day 2: *F*(1,48)=3.56, p=0.065, partial η2=0.069, BF_Inclusion_=1.05] or Group × Condition interaction [Learning Day 1: *F*(1,48)=0.749, p=0.391, partial η2=0.015, BF_Inclusion_=0.381; Recognition Day 2: *F*(1,48)=0.222, p=0.639, partial η2=0.005, BF_Inclusion_=0.313] was found for the learning scores of Day 1 or the recognition rate after 24 hours. This shows that the new group of participants, during the placebo, learned and remembered words from the M+ condition and correctly identified M-words (i.e., no meaning ascribed) at the same rate as in our previous experiment.

We then focused our analyses on learned (on Day 1) and still remembered (on Day 2) M+ new-words. In our previous work (Ripollés et al., 2016) this was the condition associated to the largest fMRI activity within the SN/VTA-HP loop, the largest physiological response and the highest subjective pleasantness ratings, even when compared with learned words that were forgotten after 24 hours (as a control, we used M-new-words correctly identified during the encoding phase and after 24 hours). Accordingly, in the present study subjective pleasantness and confidence ratings on Day 1 were higher for remembered than for forgotten M+ new-words in the 24-hour recognition test [pleasantness, *t*(25)=4.56, p<0.001, d=0.68, BF_10_=4.39; confidence, *t*(25)=2.75, p=0.011, d=0.42, BF_10_=232.82], while no difference in arousal ratings was encountered [*t*(25)=0.20, p=0.835, d=0.025, BF_10_=0.21]. In replicating our previous results (Ripollés et al., 2016), these findings confirm that intrinsic reward (i.e., derived from an internal monitoring of learning success) had a modulatory effect on long-term memory. Regarding the M-control condition, as expected, there was no difference in subjective pleasantness [*t*(25)=1.40, p=0.172, d=0.26, BF_10_=0.49], arousal [*t*(25)=1.28, p=0.212, d=0.20, BF_10_=0.43] or confidence ratings [*t*(25)=1.18, p=0.247, d=0.18, BF_10_=0.38] for M-new-words which were correctly identified during the encoding phase and still correctly rejected in the 24-hour test and those which were not. When submitting these ratings to a mixed repeated measures ANOVA with Condition (M+,M-) and Group (Pharmacological, Previous Data) as factors (in our previous analyses we excluded 4 participants from the rating analyses, see Ripollés et al., 2016; thus in this ANOVA we compare 20 participants from the previous data against 26 for the placebo session), no significant effect of Group [Pleasantness: *F*(1,44)=0.143, p=0.707, partial η2=0.003, BF_Inclusion_=0.349; Arousal: *F*(1,44)=1.66, p=0.204, partial η2=0.036, BF_Inclusion_=0.80; Confidence: *F*(1,44)=3.49, p=0.068, partial η2=0.073, BF_Inclusion_=1.17] or Group × Condition interaction [Pleasantness: *F*(1,44)=0.239, p=0.627, partial η2=0.005, BF_Inclusion_=0.321; Arousal: *F*(1,44)=0.216, p=0.645, partial η2=0.005, BF_Inclusion_=0.304; Confidence: *F*(1,44)=0.028, p=0.868, partial η2=0.001, BF_Inclusion_=0.280] was found. This shows that participants’ ratings were also in line with those of our previous experiment (Ripollés et al., 2016).

Hence, we calculated the drug effect on the behavioral data. Specifically, for the levodopa and risperidone sessions and for each subject, we calculated the percentages of change in learning scores and behavioral ratings with respect to the placebo session (see Materials and Methods). Notably, for the M+ condition, our findings show a pharmacological modulation of learning performances and behavioral reward ratings. The percentage of learned words during the encoding phase was higher under levodopa than under risperidone [as compared to placebo; *t*(25)=2.72, p=0.012, d=0.56, BF_+0_=8.26]. Importantly, this effect was still present at 24 hours for the total number of remembered new-words [*t*(25)=2.10, p=0.046, d=0.45, BF_+0_=2.62; see Figure 1A]. In addition, while no significant changes were found for the arousal ratings [*t*(25)=0.31, p=0.757, d=0.049, BF_+0_=0.26], the drug effect approached significance for confidence ratings [*t*(25)=2.05, p=0.051, d=0.51, BF_+0_=2.40] and was significant for pleasantness ratings [*t*(25)=2.70, p=0.012, d=0.64, BF_+0_=7.93], where scores for remembered words at 24 hours where higher under levodopa than under risperidone (as compared to placebo; see Figure 1B). There was not, however, a significant effect of drug on the *recognition rate* [i.e., the percentage of remembered words in the recognition test of Day 2 compared to those that were learned on Day 1; *t*(25)=-0.013, p=0.989, d=0.003, BF_+0_=0.20].

**Figure 1.**
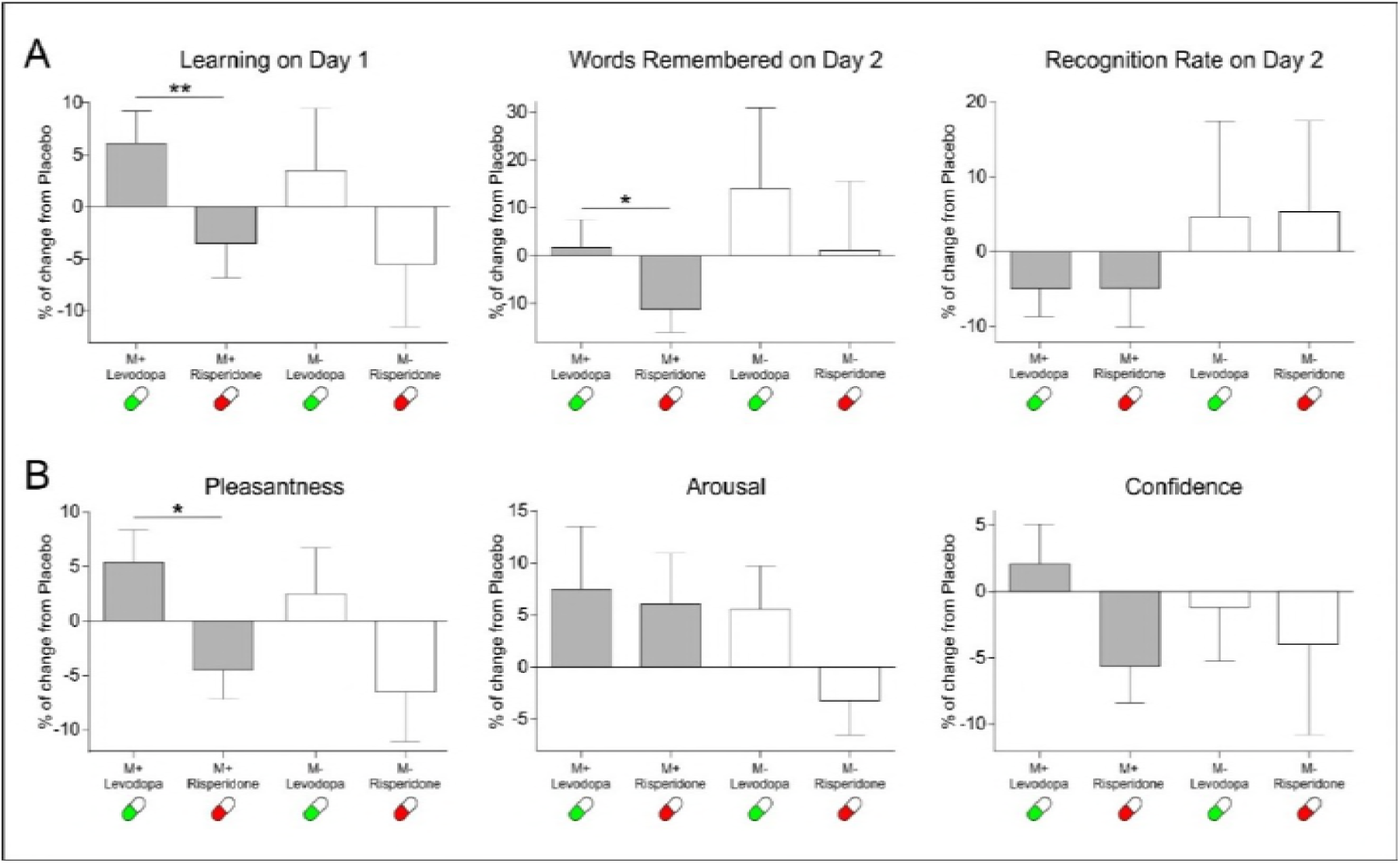
Effects of the pharmacological intervention (mean ± SEM) in. **a)** Learning scores and **b)** subjective ratings. Note that subjective ratings were only measured during the learning phase of Day 1. Effects are calculated as % of change with respect to the placebo session. *p<0.05, **p<0.01

As expected, for the control M-condition no significant differences between the risperidone and levodopa sessions as compared to placebo were found for the learning scores on Day 1 [*t*(25)=1.53, p=0.137, d=0.28, BF_+0_=1.08], the total number of correctly rejected M-words at 24 hours [*t*(25)=0.62, p=0.538, d=0.15, BF_+0_=0.35], the recognition rate [percentage of words remembered on Day 2, in respect to those learned on Day1; *t*(25)=- 0.04, p=0.968, d=0.011, BF_+0_=0.20; see Figure 1A], or the subjective ratings of arousal [*t*(23)=1.72, p=0.097, d=0.45, BF_+0_=1.46] and confidence [*t*(23)=0.36, p=0.720, d=0.10, BF_+0_=0.28; see Figure 1B; two participants were excluded from the rating analyses after not correctly rejecting any M-word at 24 hours from those correctly rejected during encoding in the levodopa session]. For the pleasantness ratings, however, the difference was close to significance [*t*(23)=2.02, p=0.055, d=0.40, BF_+0_=2.32]. However, it is important to note that the pleasantness ratings for M-trials remembered at 24 hours were *not different* from 0 at any session [Risperidone mean rating =-0.23, *t*(23)=-1.03, p=0.309, d=0.20, BF_10_=0.34; Placebo mean rating =0.24, *t*(23)=1.01, p=0.319, d=0.20, BF_10_=0.34; Levodopa mean rating =0.30, *t*(23)=1.20, p=0.240, d=0.23, BF_10_=0.40], implying that participants did not find this learning condition particularly rewarding even if the pharmacological intervention slightly modified their subjective ratings.

Given that our learning task modulates activity within the reward network (Ripollés et al., 2014 and 2016), we further tested whether individual differences in sensitivity to reward interacted with the drug intervention to modulate memory benefits (Ferreri et al., 2017; de Vries et al. 2010; Apitz and Bunzeck, 2013). Twenty-four out of the twenty-six participants completed the Physical Anhedonia Scale (PAS, Chapman et al. 1976; mean score = 11.62 ± 5.47) and we correlated (using Spearman’s rho) their individual scores with the drug effect for each learning condition (the drug effect was calculated as the subtraction of the percentage of change from placebo of the levodopa session minus the percentage of change from placebo of the risperidone session, see Materials and Methods). As a control and in order to take into account previous results (Chowdury et al., 2012), we also assessed the relationship of the learning scores with the weight dependent measure of drug dose (calculated in mg of levodopa/risperidone administered per kilogram, mean value = 1.66 ± 0.23). As expected, no significant correlations were found between the M-learning scores and the PAS [Learning Day 1 r_s_=-0.19, p = 0.372; number of correctly rejected words during Day 2, r_s_=-0.34, p=0.097; recognition rate, r_s_=-0.19, p=0.372]. In addition, no significant linear correlation or inverted U-shape relationship (Chowdury et al., 2012) was found for any learning score (M+ or M-) and the weight dependent drug dosage (all ps > 0.13). However, the drug effect for M+ trials, the number of learned words during encoding (r_s_=-0.45, p=0.025), the total number of remembered words during Day 2 (r_s_=-0.67, p<0.001) and, strikingly, the recognition rate (r_s_=-0.49, p=0.017), showed a significant correlation with the PAS (all correlations were FDR-corrected at a p<0.05 threshold, see Figure 2A; one participant was excluded from the correlations with the learning scores on Day 2 after being identified as a bivariate outlier; note that if included, the correlations become more significant: number of remembered words during Day 2, r_s_=-0.71, p<0.001; recognition rate, r_s_=-0.55, p=0.005). This suggests that the dopaminergic pharmacological intervention induced greater memory and accuracy benefits/deficits in those participants with high sensitivity to reward. Note that the drug effect for the forgetting rate, which showed no differences in memory performance when pooling all participants together, becomes significant if we divide our participants into high and low sensitivity to reward (i.e., hedonic) groups [U=35, p=0.033, η2=0.19, BF_+0_=2.81; groups divided according to the median split of their PAS scores; due to the reduced N of the groups we used a Mann-Whitney non-parametric statistical tests, see Materials and Methods and Figure 2B]. This difference was also present when comparing high/low groups for the number of words learned during encoding [U=33, p=0.024, η2=0.21, BF_+0_=4.34] and the total number of words remembered on Day 2 [U=12.5, p<0.001, η2=0.49, BF_+0_=78.74].

**Figure 2.**
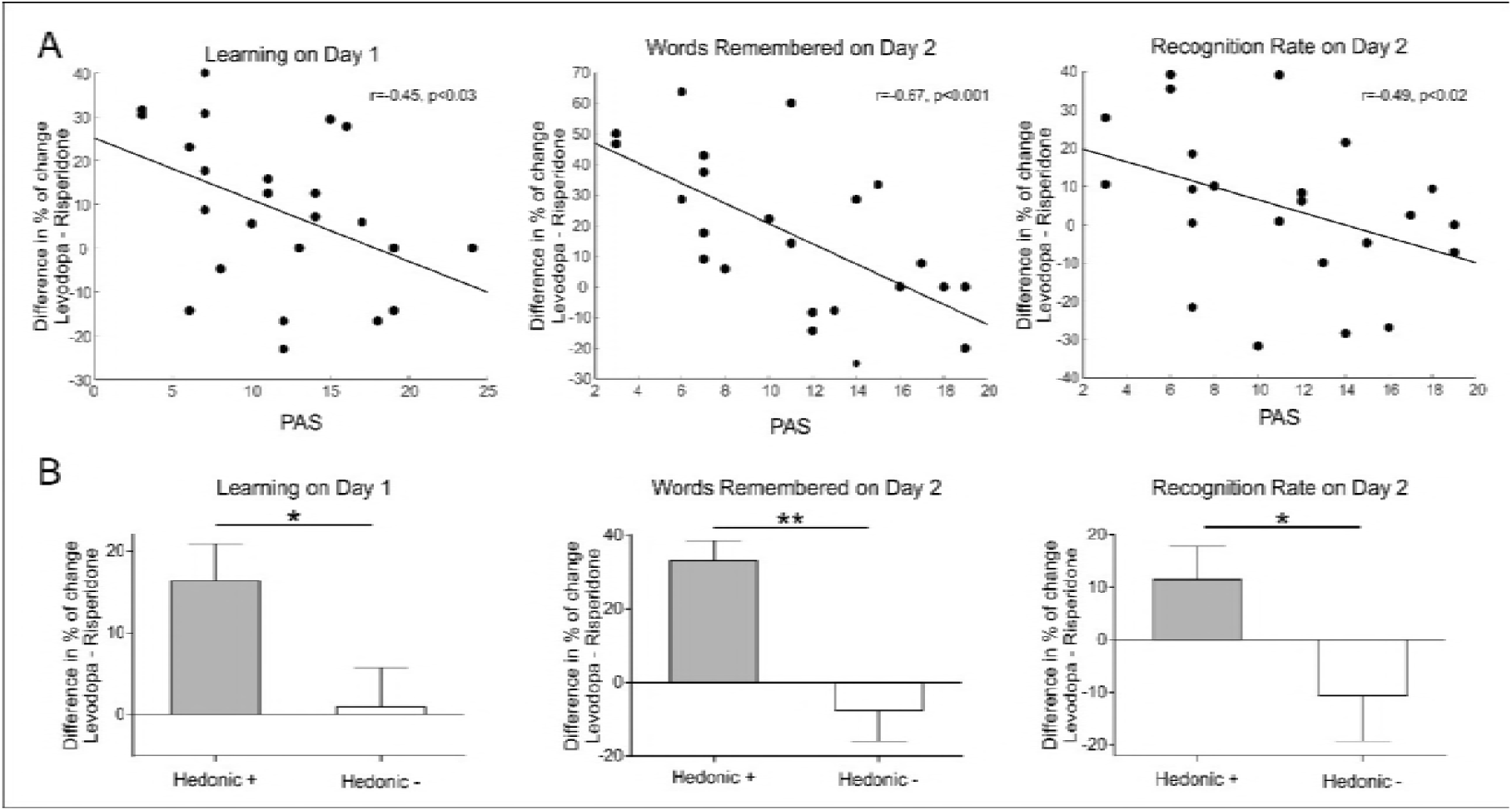
Relation between the effect of the pharmacological intervention and subjective sensitivity to reward for the learning scores. (i.e., Learning on Day 1, Words Remembered on Day 2; Recognition Rate on Day 2) obtained by **A)** correlating drug effect and PAS scores (the lower the PAS values are, the higher the general hedonia); **B)** computing the drug effect (mean ± SEM) according to high (Hedonic +) and low (Hedonic -) hedonic subjects (median split using the PAS). *p<0.05, **p<0.001.

All in all, these results show that the dopaminergic pharmacological intervention did have an effect in terms of both learning and subjective pleasantness in our learning task, inducing greater memory benefits in those participants more sensitive to reward.

## DISCUSSION

By using a double-blind, within-subject randomized pharmacological intervention during a learning task—guided by an intrinsically regulated reward process—known to activate the SN/VTA-HP loop (Ripollés et al., 2016), we showed that dopamine can modulate the entrance of new information into long-term memory. In particular, the administration of a dopaminergic precursor (levodopa) and a dopaminergic antagonist (risperidone) respectively increased and decreased both the learning rate and the level of pleasantness experienced by the participants during encoding, as well as the number of words remembered after a consolidation period (24 hours; see Figure 1B). Strikingly, the memory effects induced by the dopaminergic pharmacological intervention were stronger in participants with a higher sensitivity to reward (i.e., more hedonic; see Figure 2).

In a previous study using the same task (Ripollés et al., 2016) we showed that successful learning itself (in the absence of external feedback) was associated to increased reward processing and heightened activity within the SN/VTA and the VS. We suggested that this intrinsic reward-related signal induced a higher release of dopamine at the HP, which ultimately resulted in enhanced memory formation due to the well-known role of dopamine in mediating LTP processes. Current results provide further support for our hypothesis by showing that dopamine had an additional role during learning: participants not only learned more words (i.e., they performed better) under levodopa than under risperidone (as compared to placebo), but also found the learning experience more rewarding when stimulating, rather than blocking, the dopaminergic system. This result is in accord with previous work demonstrating that dopamine improves feedback-based learning in humans (de Vries et al., 2010) and also with research showing that internally generated signals of self-performance— driven by mesolimbic areas and in absence of external feedback—can guide and improve perceptual learning in humans (Daniel and Pollmann, 2012; Daniel and Pollmann, 2014; Guggenmos et al., 2016) and song learning (i.e., motor performance) in songbirds (Mandelblat-Cerf et al., 2014). An interesting interpretation of our results is therefore that the level of dopamine directly affected the reward value or the salience (Knetch et al., 2004) of the learning outcome in our task (i.e., learning was more enjoyable), prompting participants to be more motivated (Murty et al., 2014) and to perform better. The VS, through its connections to the prefrontal cortex (PFC; Lehericy et al., 2004; Cummings et al., 1993; Alexander et al., 1986), is located in a perfect anatomical position to add information about the relevance, salience and motivational value (Berridge and Kringelbach, 2008) of the stimuli to be learned into the SN/VTA-HP loop. As both the VS and the PFC are known to contain and receive dopaminergic receptors and projections (Haber and Knutson, 2010), dopamine might be able to alter this input, thus modulating the perceived reward underpinning the learning processes. An alternative explanation, which cannot be fully ruled out, is that the benefit in performance was driven by the suggested role of dopamine in working memory and attention (Surmeier, 2007; Brozoski et al., 1979; Linssen et al., 2014; Drijgers et al., 2012; Mehta et al., 2006). However, the fact that no significant learning benefits were induced in the control M-condition and, especially, the relationship between the learning improvements during the encoding phase and the participants’ sensitivity to reward (for a similar effect, see Ferreri et al., 2017), suggest that the learning benefit was partially driven by reward-related and dopamine-dependent processes (Diehl et al., 1992; Nieoullon et al., 2003; de Vries et al., 2010).

Previous studies in healthy humans using classical associative learning tasks (i.e., that do not usually trigger reward-related signals) have shown that levodopa intake can lead to long-term memory benefits, possibly due to the increase of the levels of dopamine in the HP (Shellshear et al., 2015; Knecht et al., 2004). However, the lack of a clear and significant memory enhancement for the control M-condition and the fact that more hedonic participants benefitted the most from the dopaminergic intervention only in the learning condition related to reward (M+), draw a more complex and perhaps more informative picture: when using a reward-based learning task (Apitz and Bunzeck, 2013; Patil et al., 2016;Oyarzun et al., 2016; Kizilirmak et al., 2015; de Vries et al., 2010), the level of memory enhancement depends on dopamine synaptic availability, but also on the individual differences in sensitivity to reward (Ferreri et al., 2017; Mas-Herrero et al., 2014; Camara et al., 2010; Marco-Pallares et al., 2009). This discovery can be crucial for dopamine-related pharmacological interventions in, for example, clinical populations with language deficits (Berthier et al., 2011). Indeed, studies with levodopa in aphasia recovery, have resulted in both positive (Seniow et al., 2009) and negative (Breitenstein et al., 2015; Leemann et al., 2011) effects. In this type of therapy, in which patients try to learn of re-learn words that are no longer accessible (Brady et al., 2012), the intensity of the language training is usually related with recovery (Bhogal et al., 2003) and it has been suggested that high training intensity may cause a ceiling effect that prevents levodopa from providing additional memory benefits (Breitenstein et al., 2015; Leemann et al., 2011). A reward-based learning task such as the one used here, along with a better understanding of the interaction between the dopaminergic precursor and the patient’s hedonic state could aid to achieve a more personalized and efficient rehabilitation success, without the need for high intensive training.

In conclusion, here we show that a dopaminergic pharmacological intervention is able to modulate behavioral measures of pleasantness, task-performance and long-term memory according to inter-individual differences in reward sensitivity. These findings further advance the idea that learning—even when achieved using a task guided by intrinsic reward—is a dopamine-dependent process, and shed new light on possible reward-based interventions for learning stimulation and/or rehabilitation.

## MATERIALS AND METHODS

### Participants

Around 150 individuals responded to advertisements and were contacted for a first phone pre-screening. Of those, 45 confirmed their availability and, after giving informed consent, were admitted at the hospital for further screening, medical examination and laboratory exams (blood and urinalysis). The study was approved by the Ethics Committee of Hospital de la Santa Creu i Sant Pau and the Spanish Medicines and Medical Devices Agency (EudraCT 2016-000801-35). The study was carried out in accordance with the Declaration of Helsinki and the ICH Good Clinical Practice Guidelines. All volunteers gave their written informed consent to participation prior to any procedure.

Subjects were judged healthy at screening 3 weeks before the first dose based on medical history, physical examination, vital signs, electrocardiogram, laboratory assessments, negative urine drug screens, and negative hepatitis B and C, and HIV serologies.

The volunteers were excluded if they had used any prescription or over-the-counter medications in the 14 days before screening, if they had a medical history of alcohol and/or drug abuse, a consumption of more than 24 or 40 grams of alcohol per day for female and male, respectively if they smoked more than 10 cigarettes/day. Women with a positive pregnancy test or not using efficient contraception methods and subjects with musical training or those unable to understand the nature and consequences of the trial or the testing procedures involved were also excluded. Additionally, volunteers were requested to abstain from alcohol, tobacco and caffeinated drinks at least during the 24 h prior to each experimental period.

Twenty nine volunteers were randomized and completed the study (19 females, mean age=22.83±4.39) in exchange of a monetary compensation according to the Spanish Legislation. The original sample size was chosen to be 30 participants, but one participant dropped out early in the study and only 29 finalized it. This sample size was selected based on several criteria, including the recommendation that, in order to achieve 80% of power, at least 30 participants should be included in an experiment in which the expected effect size is medium to large (Cohen, 1988). In addition, we took into account the sample sizes of previous studies using levodopa to modulate memory (range: between 10 and 30 participants; Apitz and Bunzeck, 2013, Copland et al., 2009, De Vries et al., 2010, Knecht et al., 2004; Chowdhury et al. 2012; Shellshear et al., 2015) and our previous behavioural studies using the same learning task (24 participants; Ripollés et al., 2016). We also computed a sample size analysis using the G*Power program, which showed that a sample size of 28 was required to ensure 80% of power to detect a significant effect (0.25) in a repeated-measures ANOVA with three sessions at the 5% significance level. We excluded 3 participants from the analyses after showing very poor memory performance on the word learning task during the placebo session (on Day 2, they remembered less than four of the M+ words learned during the encoding session). The final sample analysed for this learning paradigm consisted of 26 participants (17 women, mean age=22.27 ± 3.69).

### Study design and procedure

This double-blind, crossover, treatment sequence-randomized study was performed at the Neuropsychopharmacology Unit and Center for Drug Research (CIM) of the Santa Creu i Sant Pau Hospital of Barcelona (Spain). Experimental testing took place over three sessions. For each session, participants arrived at the hospital under fasting conditions and were given a light breakfast. Subsequently, they received in a double-blind masked fashion a capsule containing the treatment: a dopaminergic precursor with an inhibitor of peripheral dopamine metabolism (levodopa, 100 mg + carbidopa, 25 mg), a dopamine receptor antagonist (risperidone, 2mg), or placebo (lactose). After one hour of completing several behavioral tasks not described in the current manuscript, the participants completed our word learning task which lasted 45 minutes approximately. Next, participants spent their time in a resting room and were allowed to leave the hospital after 6 hours from the treatment administration. For each session, each participant came back 24 hours after for a behavioral retesting (without any pharmacological intervention), which lasted about 15 minutes. At least one week passed between one session and the other.

### Experimental word learning task

The task was virtually identical to that of our previous work (Ripollés et al., 2014, 2016 and 2017). Stimuli were presented using the Psychophysics Toolbox 3.09 (Brainard, 1997) and Matlab version R2012b. Stimuli consisted of 168 pairs of 8 word-long Spanish sentences ending in a new-word, built with an increasing degree of contextual constraint (Mestres-Missé et al., 2009; Mestres-Missé et al., 2014). Mean cloze probability (the proportion of people who complete a particular sentence fragment with a particular word) was 29.16 ± 18.95 % for the first sentence (low constraint), and 81.67 ± 11.80 % for the second (high constraint). The new-words respected the phonotactic rules of Spanish, were built by changing one or two letters of an existing word (mean number of letters= 6.02 ± 0.99) and always stood for a noun (mean frequency 43.26 ± 78.94 per million).

For each of the three different sessions, only half of the pairs of sentences disambiguated multiple meanings, thus enabling the extraction of a correct meaning for the new-word (M+ condition; e.g., 1. ‘‘Every Sunday the grandmother went to the *jedin*’’ 2. ‘‘The man was buried in the *jedin*’’; *jedin* means graveyard and is congruent with both the first and second sentence). For the other pairs, second sentences were scrambled so that they no longer matched their original first sentence. In this case, the new-word was not associated with a congruent meaning across the sentences (M-condition; e.g., 1. ‘‘Every night the astronomer watched the *heutil*’’. Moon is one possible meaning of *heutil*. 2. ‘‘In the morning break co-workers drink *heutil*.’’ Coffee is now one of the possible meanings of *heutil*, which is not congruent with the first sentence). These constituted the M-condition in which congruent meaning extraction was not possible. To ensure that both stimulus types were equally comparable, participants were told that it was just as crucial to learn the words of the M+ condition as it was to correctly reject the new-words from the M-condition.

Given that the pharmacological intervention included three sessions, we created three versions of our task that only differed in the stimuli being presented. Thus, the 168 pairs of sentences were divided into six lists of 28 pairs (as aforementioned, two conditions, M+ and M-, were presented in each of the three sessions). The six lists were created so that there were no differences (one-way ANOVA) in the cloze probability of the sentences [first sentences: *F*(5,162)=0.688, p=0.633, η2=0.021, BF_10_=0.044; second sentences: *F*(5,162)=0.419, p=0.835, η2=0.013, BF_10_=0.03], the frequency of the meanings of the real words to be learned [*F*(5,162)=1.324, p=0.256, η2=0.039, BF_10_=0.13] or the total number of letters of the new-words [*F*(5,162)=1.10, p=0.360, η2=0.033, BF_10_=0.09]. The six lists of sentences were randomly assigned in pairs to the three different sessions. Presentation of the lists was counterbalanced across the experiments so that half of the times one list was used for the M+ condition and the other half for the M-. For each participant, new-words were randomly assigned to each pair of sentences.

During each session, four pairs of M+ and four pairs of M-sentences were presented per learning block (7 blocks in total). Therefore, a total of 28 new-words from the M+ and 28 from the M-conditions were presented during each of the three sessions. In order to achieve an ecologically valid paradigm, presentation of the first and second sentences with the same new-word at the end were separated in time. The 4 first sentences of each of the M+ and M-conditions (a total of eight new-words) were presented in a pseudo-randomized order (e.g., M+1A, M-1A, M-1B, M-1C, M+1B, M+1C, M+1D, M-1D). Then, the second ‘pair’ sentences of both M+ and M-conditions were presented (i.e., second presentation of the identical eight new-words), again in a pseudo-randomized order (e.g., M-2C, M-2B, M+2B, M+2D, M-2D, M+2C, M+2A, M-2A). The temporal order of the different new-words during first sentence presentation was not related in any systematic way to the order of presentation of the same new-words for their second sentence. Participants were instructed to produce a verbal answer 8 seconds after the new-word of a second sentence appeared. If participants thought that the new-word had a congruent meaning, they had to provide its meaning in Spanish (e.g., *graveyard*). If the new-word had no consistent meaning, they had to say the word *incongruent.* If they did not know whether the new-word had a consistent meaning or not, they had to remain silent. Vocal answers were recorded and later corrected (for the M+ condition, incorrect answers included misses, providing the wrong meaning or saying *incongruent;* for the M-condition, incorrect answers included misses or providing any meaning at all). After giving a verbal answer, participants first provided a confidence rating that allowed for the assessment of the subjective evaluation of their performance. Specifically, subjects were requested to enter, using the keyboard, a value between -4 and 4 (9 point scale with 0 as the neutral value). Then, participants had to rate their emotions with respect to arousal and pleasantness using the 9-point (as with confidence ratings, from -4 to 4) visual *Self-Assessment Manikin* scale (SAM). For valence/pleasantness, the SAM ranges from a sad, frowning figure (i.e., very negative) to a happy, smiling figure (i.e., very positive). For arousal, the SAM ranges from a relaxed figure (i.e., very calm) to an excited figure (i.e., very aroused). All participants completed a training block to familiarize them with the task.

Each trial started with a fixation cross lasting 1000 ms, continued with the 7 first Spanish words of the sentence presented for 2 seconds, and was followed by a 1 second duration dark screen. The new-word was presented for 1000 ms. and was followed by 7 seconds of a small fixation point presented in the middle of the screen. For first sentences, a new trial was presented after 3 seconds of dark screen. For second sentences, after this period, a screen with the word *Answer* appeared and subjects had 3 seconds to produce a verbal answer. Then, the confidence and SAM scales for pleasantness and arousal were sequentially presented (the experiment did not continue until participants provided a rating). Finally, a new second sentence trial started after 3 seconds of dark screen. All words were placed in the middle of a black screen with a font size of 22 and in white color.

To avoid biasing our results, participants were not told at any point prior to the start of the experiment that the goal of the study was to assess whether the learning of a new-word and its meaning was intrinsically rewarding. Instead, they were told that the objective of the study was to assess how reading load affects mood and that, in order to ensure that there was a real reading load, they had to learn the words of the M+ condition and to detect the incongruence of the new-words from the M-. Finally, participants were told that they had to give pleasantness and arousal ratings when the second sentences appeared because that moment signaled that reading load had already occurred (i.e., half of the encoding block had already elapsed). After the experiment, participants were first questioned about the objective of the study. None of them answered that it was to assess whether word-learning was rewarding.

Approximately 24 hours after the learning lesson ended, participants returned to the lab to complete a recognition test (note that no drugs were administered to subjects on Day 2). In this test, participants were presented, in a pseudo-randomized order, with all the 28 M+ and 28 M-new-words used during the encoding session. This test was devised in order to assess which of the learned words during encoding were still remembered and which of them had been forgotten after a 24 hour retention period. Participants were aware that they would complete this test before completing the encoding session. It was made explicit that they would assess both M+ and M-new-words during the test phase. In the test, participants were presented with a new-word at the centre of the screen with two possible meanings below: one on the left and one on the right. If the new-word tested did not have a congruent meaning associated between the first and the second sentence, and thus correct meaning extraction was not possible (M-condition), participants had to press a button located in their left hand. In this case, the two possible meanings presented served as foils: one was the meaning evoked by the second sentence of the M-new-word being tested; the other word shown was the meaning evoked by another second sentence presented in the same run as the new-word being tested. Instead, if the new-word tested had a consistent meaning through the first and second sentence, and thus correct meaning extraction was possible (M+ condition), participants had to select the correct meaning among the two presented. In this case, one of the two possible meanings was correct and the other, which served as a foil, was the meaning of another new-word presented in the same run. In addition, participants could also press a fourth button if they did not know the answer. Thus chance level was at 25% (no consistent meaning, consistent meaning on the left, consistent meaning on the right, not remembered).

### Statistical Analyses for confidence, pleasantness and arousal subjective scales and learning scores for encoding and retrieval

We first assessed whether the results of the placebo session replicated our previous behavioral data (Experiment 3 in Ripollés et al., 2016). Besides the three subjective ratings, for these first comparisons, we used two learning scores: the percentage of words learned on Day 1 (total number of words learned divided by the total number of words presented) and the recognition rate (total number of words learned during encoding and remembered on Day 2 divided by the number of words learned during Day 1). For the analyses regarding the subjective scales, we divided our M+ trials into those in which subjects learned the new-word during the learning session and still remembered it in the test after the recognition test (*remembered* condition) and those in which the new-word was not correctly identified in the 24 hour test (*forgotten* condition). We used the same approach to divide the M-trials into those in which a word was correctly marked as incongruent during encoding and still correctly rejected after 24 hours and those in which the new-word was not correctly rejected in the follow-up test. To replicate our previous results, we first used paired t-tests to compare whether ratings for confidence, arousal and pleasantness were greater for remembered than for forgotten M+ and M-new-words. We then submitted both the ratings and the learning scores to a mixed repeated measures ANOVA with Condition (M+,M-) as a within-subjects variable and Group (Pharmacological Group, Exp. 3 in Ripollés et al., 2016) as a between subjects variable.

Given that current behavioral results replicate our previous work (see results) and that in our previous study (Ripollés et al., 2016) remembered M+ words were the trials showing the highest fMRI activity within the SN/VTA-HP loop, the largest physiological response and the highest subjective pleasantness ratings, we focused all the analyses regarding the effect of the pharmacological intervention in the trials in which a word was learned during encoding and still remembered during the recognition test at 24 hours (M+ condition). For the control condition, we used those M-trials in which a word was correctly rejected during both encoding and the follow-up test. As measures for memory effects, we used the total number of words learned during encoding and remembered on the follow-up test and the percentage remembered words in the recognition test compared to the number of learned words during the learning phase (i.e., the recognition rate). In order to control for individual differences, we used the placebo session as a baseline. Thus, for each learning score and subjective scale we calculated the percentage of change from the placebo session [e.g., (levodopa score - placebo score)/(placebo score)]. Therefore, for each participant, learning score and subjective scale, we obtained the percentage of change from placebo of the risperidone and levodopa sessions. We used paired t-tests to calculate whether the difference between the changes induced by the risperidone and levodopa sessions were significant.

For the correlations between the learning scores and the PAS we used Spearman’s rho with a p<0.05 FDR correction to account for the 3 different correlations calculated per condition. The PAS was used as a proxy to reflect the degree of pleasure taken by individuals when engaging in rewarding behavior (Der-Avakian et al., 2012). Note that two participants were excluded from this analysis as they did not complete the PAS. We also correlated the learning scores with a weight dependent measure of drug dose, calculated in mg of levodopa/risperidone per kilogram. Finally, we used the median PAS value to split our final sample of 24 participants into high and low hedonic groups. For the learning scores, we first calculated the drug effect as a subtraction of the percentage of change from placebo induced by the levodopa session minus that induced by the risperidone session. We then assessed were the total drug effect for the learning scores was different for high vs. low hedonic groups by using a non-parametric test Mann-Whitney (to better account for the reduced number of participants in each group).

For significant interactions of mixed between-within ANOVA models, partial eta squares (η2) is provided as a measure of effect size. For significant differences in between group one-way ANOVAs, eta squares (η2) is provided (calculated by dividing the between groups sum of squares by the total sum of squares). For significant differences measured with t-tests, Cohen’s d is provided after applying Hedges’ correction (the average of the standard deviation of the variables being compared was used as a standardizer; Cumming, 2012). For significant differences measured with the Mann-Whitney test, eta squares (η2) is provided (calculated as Z^2^/N)

In addition, confirmatory Bayesian statistical analyses were computed with the software JASP using default priors (JASP Team, 2018; Morey and Rouder, 2015; Rouder and Morey, 2012; Wagenmakers et al., 2018b; Wagenmakers et al., 2018a). We reported Bayes factors (BF_10_), which reflect how likely data is to arise from one model, compared, in our case, to the null model (i.e., the probability of the data given H1 relative to H0). For comparisons with a strong a priori, the alternative hypothesis was specified so that one group/condition was greater than the other (BF_+0_). We did this, specifically, for the drug effects comparisons in which we expected levodopa and risperidone to facilitate and disrupt learning/ratings, respectively; and for the group comparisons in which we expected more hedonic participants to remember more words than less hedonic participants. For mixed within-between models we used the Bayes Inclusion factor based on matched models, representing the evidence for all models containing a particular effect to equivalent models stripped of that effect (BF_Inlcusion_, also called Baws factor).

## ACKNOWLEDGMENTS

We thank the staff of the Centre d’Investigació del Medicament de l’Institut de Recerca HSCSP for their help. The present project has been funded by the Spanish Government (MINECO Grant PSI2011-29219 to A.R.F. and AP2010-4179 to P.R.) and German Research Council (DFG-SFB-779/A15 to T.N.). L.F. was supported by Morelli-Rotary postdoctoral fellowship. M.V. was partially supported by FIS trough grant CP04/00 121 from the Spanish Health Ministry in collaboration with Institut de Recerca de l’Hospital de la Santa Creu i Sant Pau, Barcelona; she is a member of CIBERSAM (funded by the Spanish Health Ministry, Instituto de Salud Carlos III).

## ADDITIONAL INFORMATION

### Author contributions

PR, LF, JR & ARF designed research; LF, EMH, HA, AGA, RA and MV performed research; PR, LF, EMH, JMP and TN analyzed data; All authors participated in the drafting of the manuscript.

### Ethics

This study was performed according to local ethics and to the Declaration of Helsinki. It was approved by the Ethics Committee of Hospital Sant Pau and by the Spanish Medicines and Medical Devices Agency (EudraCT 2016-000801-35). All participants gave informed written consent and received compensation for their participation in the study according to Spanish legislation.

## REFERENCES

Adcock, R.A., Thangavel, A., Whitfield-Gabrieli, S., Knutson, B., & Gabrieli, J.D. 2006. Reward-motivated learning: mesolimbic activation precedes memory formation. Neuron, 50, (3) 507–517 available from: PM:16675403

Alexander, G.E., DeLong, M.R., & Strick, P.L. 1986. Parallel organization of functionally segregated circuits linking basal ganglia and cortex. Annu.Rev.Neurosci., 9, 357–381 available from: PM:3085570

Apitz, T. & Bunzeck, N. 2013. Dopamine controls the neural dynamics of memory signals and retrieval accuracy. Neuropsychopharmacology, 38, (12) 2409–2417 available from: PM:23728140

Berridge, K.C. & Kringelbach, M.L. 2008. Affective neuroscience of pleasure: reward in humans and animals. Psychopharmacology, 199, (3) 457–480 available from: PM:18311558

Berthier, M.L., Pulvermuller, F., Davila, G., Casares, N.G., & Gutierrez, A. 2011. Drug therapy of post-stroke aphasia: a review of current evidence. Neuropsychol.Rev., 21, (3) 302–317 available from: PM:21845354

Bethus, I., Tse, D., & Morris, R.G. 2010. Dopamine and memory: modulation of the persistence of memory for novel hippocampal NMDA receptor-dependent paired associates. J.Neurosci., 30, (5) 1610–1618 available from: PM:20130171

Bhogal, S.K., Teasell, R., & Speechley, M. 2003. Intensity of aphasia therapy, impact on recovery. Stroke, 34, (4) 987–993 available from: PM:12649521

Boehler, C.N., Hopf, J.M., Krebs, R.M., Stoppel, C.M., Schoenfeld, M.A., Heinze, H.J., & Noesselt, T. 2011. Task-load-dependent activation of dopaminergic midbrain areas in the absence of reward. J.Neurosci., 31, (13) 4955–4961 available from: PM:21451034

Brady, M.C., Kelly, H., Godwin, J., & Enderby, P. 2012. Speech and language therapy for aphasia following stroke. Cochrane.Database.Syst.Rev., 5, CD000425 available from: PM:22592672

Brainard, D.H. 1997. The Psychophysics Toolbox. Spat.Vis., 10, (4) 433–436 available from: PM:9176952

Breitenstein, C., Korsukewitz, C., Baumgartner, A., Floel, A., Zwitserlood, P., Dobel, C., & Knecht, S. 2015. L-dopa does not add to the success of high-intensity language training in aphasia. Restor.Neurol.Neurosci., 33, (2) 115–120 available from: PM:25588456

Breitenstein, C., Wailke, S., Bushuven, S., Kamping, S., Zwitserlood, P., Ringelstein, E.B., & Knecht, S. 2004. D-amphetamine boosts language learning independent of its cardiovascular and motor arousing effects. Neuropsychopharmacology, 29, (9) 1704–1714 available from: PM:15114342

Brozoski, T.J., Brown, R.M., Rosvold, H.E., & Goldman, P.S. 1979. Cognitive deficit caused by regional depletion of dopamine in prefrontal cortex of rhesus monkey. Science, 205, (4409) 929–932 available from: PM:112679

Bunzeck, N. & Duzel, E. 2006. Absolute coding of stimulus novelty in the human substantia nigra/VTA. Neuron, 51, (3) 369–379 available from: PM:16880131

Bunzeck, N., Guitart-Masip, M., Dolan, R.J., & Duzel, E. 2014. Pharmacological dissociation of novelty responses in the human brain. Cereb.Cortex, 24, (5) 1351–1360 available from: PM:23307638

Callan, D.E. & Schweighofer, N. 2008. Positive and negative modulation of word learning by reward anticipation. Hum.Brain Mapp., 29, (2) 237–249 available from: PM:17390317

Camara, E., Kramer, U.M., Cunillera, T., Marco-Pallares, J., Cucurell, D., Nager, W., Mestres-Misse, A., Bauer, P., Schule, R., Schols, L., Tempelmann, C., Rodriguez-Fornells, A., & Munte, T.F. 2010. The effects of COMT (Val108/158Met) and DRD4 (SNP -521) dopamine genotypes on brain activations related to valence and magnitude of rewards. Cereb.Cortex, 20, (8) 1985–1996 available from: PM:20038544

Chapman, L.J., Chapman, J.P., & Raulin, M.L. 1976. Scales for physical and social anhedonia. J.Abnorm.Psychol, 85, (4) 374–382 available from: PM:956504

Chowdhury, R., Guitart-Masip, M., Bunzeck, N., Dolan, R.J., & Duzel, E. 2012. Dopamine modulates episodic memory persistence in old age. J.Neurosci., 32, (41) 14193–14204 available from: PM:23055489

Cohen, J. 1988. Statistical Power Analysis for the Behavioral Sciences Hillsdale, New Jersey, Lawrence Erlbaum Associates.

Copland, D.A., McMahon, K.L., Silburn, P.A., & de Zubicaray, G.I. 2009. Dopaminergic neuromodulation of semantic processing: a 4-T FMRI study with levodopa. Cereb.Cortex, 19, (11) 2651–2658 available from: PM:19321651

Cummings, J.L. 1993. Frontal-subcortical circuits and human behavior. Arch.Neurol., 50, (8) 873–880 available from: PM:8352676

Daniel, R. & Pollmann, S. 2012. Striatal activations signal prediction errors on confidence in the absence of external feedback. Neuroimage., 59, (4) 3457–3467 available from: PM:22146752

Daniel, R. & Pollmann, S. 2014. A universal role of the ventral striatum in reward-based learning: evidence from human studies. Neurobiol.Learn.Mem., 114, 90–100 available from: PM:24825620

de Vries, M.H., Ulte, C., Zwitserlood, P., Szymanski, B., & Knecht, S. 2010. Increasing dopamine levels in the brain improves feedback-based procedural learning in healthy participants: an artificial-grammar-learning experiment. Neuropsychologia, 48, (11) 3193–3197 available from: PM:20600185

Der-Avakian, A. & Markou, A. 2012. The neurobiology of anhedonia and other reward-related deficits. Trends Neurosci., 35, (1) 68–77 available from: PM:22177980

Diehl, D.J. & Gershon, S. 1992. The role of dopamine in mood disorders. Compr.Psychiatry, 33, (2) 115–120 available from: PM:1347497

Drijgers, R.L., Verhey, F.R., Tissingh, G., van Domburg, P.H., Aalten, P., & Leentjens, A.F. 2012. The role of the dopaminergic system in mood, motivation and cognition in Parkinson’s disease: a double blind randomized placebo-controlled experimental challenge with pramipexole and methylphenidate. J.Neurol.Sci., 320, (1-2) 121–126 available from: PM:22824349

Duzel, E., Bunzeck, N., Guitart-Masip, M., Wittmann, B., Schott, B.H., & Tobler, P.N. 2009. Functional imaging of the human dopaminergic midbrain. Trends Neurosci., 32, (6) 321–328 available from: PM:19446348

Ferenczi, E.A., Zalocusky, K.A., Liston, C., Grosenick, L., Warden, M.R., Amatya, D., Katovich, K., Mehta, H., Patenaude, B., Ramakrishnan, C., Kalanithi, P., Etkin, A., Knutson, B., Glover, G.H., & Deisseroth, K. 2016. Prefrontal cortical regulation of brainwide circuit dynamics and reward-related behavior. Science, 351, (6268) aac9698 available from: PM:26722001

Ferreri, L. & Rodriguez-Fornells, A. 2017. Music-related reward responses predict episodic memory performance. Exp.Brain Res. available from: PM:28940086

Frey, U., Schroeder, H., & Matthies, H. 1990. Dopaminergic antagonists prevent long-term maintenance of posttetanic LTP in the CA1 region of rat hippocampal slices. Brain Res., 522, (1) 69–75 available from: PM:1977494

Goto, Y. & Grace, A.A. 2005. Dopaminergic modulation of limbic and cortical drive of nucleus accumbens in goal-directed behavior. Nat.Neurosci., 8, (6) 805–812 available from: PM:15908948

Guggenmos, M., Wilbertz, G., Hebart, M.N., & Sterzer, P. 2016. Mesolimbic confidence signals guide perceptual learning in the absence of external feedback. Elife., 5, available from: PM:27021283

Guitart-Masip, M., Bunzeck, N., Stephan, K.E., Dolan, R.J., & Duzel, E. 2010. Contextual novelty changes reward representations in the striatum. J.Neurosci., 30, (5) 1721–1726 available from: PM:20130181

Haber, S.N. & Knutson, B. 2010. The reward circuit: linking primate anatomy and human imaging. Neuropsychopharmacology, 35, (1) 4–26 available from: PM:19812543

Hansen, N. & Manahan-Vaughan, D. 2014. Dopamine D1/D5 receptors mediate informational saliency that promotes persistent hippocampal long-term plasticity. Cereb.Cortex, 24, (4) 845–858 available from: PM:23183712

Huang, Y.Y. & Kandel, E.R. 1995. D1/D5 receptor agonists induce a protein synthesis-dependent late potentiation in the CA1 region of the hippocampus. Proc.Natl.Acad.Sci.U.S.A, 92, (7) 2446–2450 available from: PM:7708662

JASP Team (2018) JASP (Version 0.8.6) [Computer software].

Kaminski, J., Mamelak, A.N., Birch, K., Mosher, C.P., Tagliati, M., & Rutishauser, U. 2018. Novelty-Sensitive Dopaminergic Neurons in the Human Substantia Nigra Predict Success of Declarative Memory Formation. Curr.Biol. available from: PM:29657115

Kizilirmak, J.M., Galvao Gomes da, S.J., Imamoglu, F., & Richardson-Klavehn, A. 2015. Generation and the subjective feeling of “aha!” are independently related to learning from insight. Psychol.Res. available from: PM:26280758

Knecht, S., Breitenstein, C., Bushuven, S., Wailke, S., Kamping, S., Floel, A., Zwitserlood, P., & Ringelstein, E.B. 2004. Levodopa: faster and better word learning in normal humans. Ann.Neurol., 56, (1) 20–26 available from: PM:15236398

Knutson, B. & Gibbs, S.E. 2007. Linking nucleus accumbens dopamine and blood oxygenation. Psychopharmacology (Berl), 191, (3) 813–822 available from: PM:17279377

Leemann, B., Laganaro, M., Chetelat-Mabillard, D., & Schnider, A. 2011. Crossover trial of subacute computerized aphasia therapy for anomia with the addition of either levodopa or placebo. Neurorehabil.Neural Repair, 25, (1) 43–47 available from: PM:20834044

Lehericy, S., Ducros, M., Van de Moortele, P.F., Francois, C., Thivard, L., Poupon, C., Swindale, N., Ugurbil, K., & Kim, D.S. 2004. Diffusion tensor fiber tracking shows distinct corticostriatal circuits in humans. Ann.Neurol., 55, (4) 522–529 available from: PM:15048891

Linssen, A.M., Sambeth, A., Vuurman, E.F., & Riedel, W.J. 2014. Cognitive effects of methylphenidate in healthy volunteers: a review of single dose studies. Int.J.Neuropsychopharmacol., 17, (6) 961–977 available from: PM:24423151

Lisman, J., Grace, A.A., & Duzel, E. 2011. A neoHebbian framework for episodic memory; role of dopamine-dependent late LTP. Trends Neurosci., 34, (10) 536–547 available from: PM:21851992

Lisman, J.E. & Grace, A.A. 2005. The hippocampal-VTA loop: controlling the entry of information into long-term memory. Neuron, 46, (5) 703–713 available from: PM:15924857

Mandelblat-Cerf, Y., Las, L., Denisenko, N., & Fee, M.S. 2014. A role for descending auditory cortical projections in songbird vocal learning. Elife., 3, available from: PM:24935934

Marco-Pallares, J., Cucurell, D., Cunillera, T., Kramer, U.M., Camara, E., Nager, W., Bauer, P., Schule, R., Schols, L., Munte, T.F., & Rodriguez-Fornells, A. 2009. Genetic variability in the dopamine system (dopamine receptor D4, catechol-O-methyltransferase) modulates neurophysiological responses to gains and losses. Biol.Psychiatry, 66, (2) 154–161 available from: PM:19251248

Mas-Herrero, E., Zatorre, R.J., Rodriguez-Fornells, A., & Marco-Pallares, J. 2014. Dissociation between musical and monetary reward responses in specific musical anhedonia. Curr.Biol., 24, (6) 699–704 available from: PM:24613311

McNamara, C.G., Tejero-Cantero, A., Trouche, S., Campo-Urriza, N., & Dupret, D. 2014. Dopaminergic neurons promote hippocampal reactivation and spatial memory persistence. Nat.Neurosci., 17, (12) 1658–1660 available from: PM:25326690

Mehta, M.A. & Riedel, W.J. 2006. Dopaminergic enhancement of cognitive function. Curr.Pharm.Des, 12, (20) 2487–2500 available from: PM:16842172

Mestres-Misse, A., Munte, T.F., & Rodriguez-Fornells, A. 2009. Functional neuroanatomy of contextual acquisition of concrete and abstract words. J.Cogn Neurosci., 21, (11) 2154–2171 available from: PM:19199404

Mestres-Misse, A., Munte, T.F., & Rodriguez-Fornells, A. 2014. Mapping concrete and abstract meanings to new words using verbal contexts. Second Language Research, 30, (2) 191–223

Mestres-Misse, A., Rodriguez-Fornells, A., & Munte, T.F. 2007. Watching the brain during meaning acquisition. Cereb.Cortex, 17, (8) 1858–1866 available from: PM:17056648

Mestres-Misse, A., Rodriguez-Fornells, A., & Munte, T.F. 2010. Neural differences in the mapping of verb and noun concepts onto novel words. Neuroimage., 49, (3) 2826–2835 available from: PM:19837174

Morey RD, Rouder JN (2015) BayesFactor (Version 0.9.10-2)[Computer software]

Murty, V.P. & Adcock, R.A. 2014. Enriched encoding: reward motivation organizes cortical networks for hippocampal detection of unexpected events. Cereb.Cortex, 24, (8) 2160–2168 available from: PM:23529005

Nieoullon, A. & Coquerel, A. 2003. Dopamine: a key regulator to adapt action, emotion, motivation and cognition. Curr.Opin.Neurol., 16 Suppl 2, S3–S9 available from: PM:15129844

Oyarzun, J.P., Packard, P.A., de Diego-Balaguer, R., & Fuentemilla, L. 2016. Motivated encoding selectively promotes memory for future inconsequential semantically-related events. Neurobiol.Learn.Mem., 133, 1–6 available from: PM:27224885

Patil, A., Murty, V.P., Dunsmoor, J.E., Phelps, E.A., & Davachi, L. 2017. Reward retroactively enhances memory consolidation for related items. Learn.Mem., 24, (1) 65–69 available from: PM:27980078

Ripolles, P., Biel, D., Penaloza, C., Kaufmann, J., Marco-Pallares, J., Noesselt, T., & Rodriguez-Fornells, A. 2017. Strength of Temporal White Matter Pathways Predicts Semantic Learning. J.Neurosci., 37, (46) 11101–11113 available from: PM:29025925

Ripolles, P., Marco-Pallares, J., Alicart, H., Tempelmann, C., Rodriguez-Fornells, A., & Noesselt, T. 2016. Intrinsic monitoring of learning success facilitates memory encoding via the activation of the SN/VTA-Hippocampal loop. Elife., 5, (e17441) available from: PM:27644419

Ripolles, P., Marco-Pallares, J., Hielscher, U., Mestres-Misse, A., Tempelmann, C., Heinze, H.J., Rodriguez-Fornells, A., & Noesselt, T. 2014. The Role of Reward in Word Learning and Its Implications for Language Acquisition. Curr.Biol., 24, (21) 2606–2611 available from: PM:25447993

Rossato, J.I., Bevilaqua, L.R., Izquierdo, I., Medina, J.H., & Cammarota, M. 2009. Dopamine controls persistence of long-term memory storage. Science, 325, (5943) 1017–1020 available from: PM:19696353

Rouder JN, Morey RD (2012) Default Bayes Factors for Model Selection in Regression. Multivariate Behav Res 47:877–903.

Salimpoor, V.N., Benovoy, M., Larcher, K., Dagher, A., & Zatorre, R.J. 2011. Anatomically distinct dopamine release during anticipation and experience of peak emotion to music. Nat.Neurosci., 14, (2) 257–262 available from: PM:21217764

Schott, B.H., Minuzzi, L., Krebs, R.M., Elmenhorst, D., Lang, M., Winz, O.H., Seidenbecher, C.I., Coenen, H.H., Heinze, H.J., Zilles, K., Duzel, E., & Bauer, A. 2008. Mesolimbic functional magnetic resonance imaging activations during reward anticipation correlate with reward-related ventral striatal dopamine release. J.Neurosci., 28, (52) 14311–14319 available from: PM:19109512

Seniow, J., Litwin, M., Litwin, T., Lesniak, M., & Czlonkowska, A. 2009. New approach to the rehabilitation of post-stroke focal cognitive syndrome: effect of levodopa combined with speech and language therapy on functional recovery from aphasia. J.Neurol.Sci., 283, (1-2) 214–218 available from: PM:19268976

Shellshear, L., MacDonald, A.D., Mahoney, J., Finch, E., McMahon, K., Silburn, P., Nathan, P.J., & Copland, D.A. 2015. Levodopa enhances explicit new-word learning in healthy adults: a preliminary study. Hum.Psychopharmacol. available from: PM:25900350

Shohamy, D. & Adcock, R.A. 2010. Dopamine and adaptive memory. Trends Cogn Sci., 14, (10) 464–472 available from: PM:20829095

Surmeier, D.J. 2007. Dopamine and working memory mechanisms in prefrontal cortex. J.Physiol, 581, (Pt 3) 885 available from: PM:17495037

Wagenmakers EJ, et al. (2018a) Bayesian inference for psychology. Part II: Example applications with JASP. Psychon Bull Rev 25:58–76.

Wagenmakers EJ, Marsman M, Jamil T, Ly A, Verhagen J, Love J, Selker R, Gronau QF, Smira M, Epskamp S, Matzke D, Rouder JN, Morey RD (2018b) Bayesian inference for psychology. Part I: Theoretical advantages and practical ramifications. Psychon Bull Rev 25:35–57.

Whiting, E., Chenery, H., Chalk, J., Darnell, R., & Copland, D. 2007. Dexamphetamine enhances explicit new word learning for novel objects. Int.J.Neuropsychopharmacol., 10, (6) 805–816 available from: PM:17250775

Whiting, E., Chenery, H.J., Chalk, J., Darnell, R., & Copland, D.A. 2008. The explicit learning of new names for known objects is improved by dexamphetamine. Brain Lang, 104, (3) 254–261 available from: PM:17428528

Wittmann, B.C., Schott, B.H., Guderian, S., Frey, J.U., Heinze, H.J., & Duzel, E. 2005. Reward-related FMRI activation of dopaminergic midbrain is associated with enhanced hippocampus-dependent long-term memory formation. Neuron, 45, (3) 459–467 available from: PM:15694331

Wolosin, S.M., Zeithamova, D., & Preston, A.R. 2012. Reward modulation of hippocampal subfield activation during successful associative encoding and retrieval. J.Cogn Neurosci., 24, (7) 1532–1547 available from: PM:22524296

